# Submaximal eccentric training during immobilization does not prevent serial sarcomere loss or impairments in mechanical function in old or young rats

**DOI:** 10.1101/2024.10.18.619056

**Authors:** Avery Hinks, Ethan Vlemmix, Geoffrey A. Power

## Abstract

The age-related loss of muscle mass is partly driven by a reduction in serial sarcomere number (SSN), and further SSN loss occurs during immobilization. SSN is associated with optimal force and power production and muscle passive tension, thus immobilization-induced SSN loss is especially a concern for older individuals who are often subjected to forced muscle disuse with illness and injury. We previously showed that submaximal eccentric resistance training increased SSN and improved muscle function in old rats. The present study investigated whether this training could prevent the losses of SSN and function when performed intermittently during immobilization. 10 old (32 months) and 10 young (8 months) rats underwent unilateral casting of the plantar flexors in a shortened position for 2 weeks. Thrice weekly, casts were removed for isokinetic eccentric resistance training. Pre- and post-training we assessed *in-vivo* maximum isometric torque at ankle angles corresponding to stretched and neutral muscle lengths, the passive torque-angle relationship, and isotonic power. The soleus and medial gastrocnemius were harvested for SSN measurements, with the untrained leg as a control. In old and young rats, muscles of the casted leg had lower muscle wet weights (−20-40%), physiological cross-sectional area (−16-20%), and SSN (−7-29%) than the control leg. Furthermore, maximum isometric torque (−37-46%) and isotonic power (−≈70%) decreased, and passive torque increased (+≈400%) from pre- to post-training for both age groups. Thus, irrespective of age, submaximal eccentric resistance training 3 days/week was ineffective for preventing the losses of muscle contractile tissue and mechanical function during casting.

## Introduction

Sarcopenia, the age-related loss of muscle mass, is in part driven by the loss of sarcomeres aligned in series along a muscle ^1,2^. This age-related loss of serial sarcomere number (SSN) has been shown to be 14-35% in old compared to young rodents ^3–6^ and helps explain the 7-35% reductions in muscle fascicle length (FL) as measured by ultrasound observed in older humans ^7–17^. SSN is also associated with the width of the plateau region of the muscle force-length relationship (e.g., the range of motion in which the muscle can generate optimal force), maximum power production, and passive tension ^18^, thus the age-related loss of SSN likely contributes to impairments in these muscle mechanical properties with aging. Loss of SSN also occurs when muscle is immobilized in a shortened position ^19–22^. We recently showed in old rats that with 2 weeks of casting in full plantar flexion (i.e., shortened soleus muscle), the soleus loses an additional 25% SSN on top of that already lost with aging ^6^. Furthermore, old rats took longer to recover SSN than young rats following cast removal ^6^. The compounded losses of SSN with aging and immobilization are thereby especially a concern for older adult humans, who are prone to falls and injuries that could result in extended periods of forced muscle disuse ^23,24^.

Previously, Williams et al. (1990) showed that 0.5-2 hours of passive stretching per day during 2 weeks of casting prevented soleus SSN loss in young adult mice. However, passive stretching for 40 minutes only once ^26^ or thrice ^27^ weekly did not prevent soleus SSN loss during casting in young adult rats, suggesting only long and potentially unfeasible passive stretching interventions can prevent SSN loss. Compared to passive stretching, eccentric resistance training has shown better effectiveness for increasing SSN or FL and improving mechanical performance in both animals and humans ^28–31^, likely due to the greater mechanical tension and associated mechanotransductive response for muscle growth ^18,32^. We recently showed that 4 weeks of submaximal (60% of maximum) eccentric training for just ∼10 minutes/day 3 days/week increased SSN of the old rat plantar flexors by 8-11%, notably restoring SSN to a level similar to that of young rats ^33^. This increase in SSN was accompanied by an improvement in torque production throughout the joint range of motion, a 23% improvement in peak isotonic power, and a reduction in passive torque ^33^. Similar eccentric resistance training interventions have also yielded performance improvements in older humans ^34–36^. However, whether submaximal eccentric training 3 days/week can prevent SSN loss during immobilization in young and old rats is unknown.

The present study assessed whether submaximal eccentric training 3 days/week during two weeks of casting can prevent the losses of SSN and muscle mechanical performance in old and young rats. Due to the large benefits of submaximal eccentric training for SSN and muscle mechanical performance in old rats that we observed previously ^33^, we hypothesized that this training would ameliorate the effects of casting on morphological and mechanical function in both old and young rats.

## Methods

### Animals

10 old (32 months) and 10 young (8 months) male Fisher 344/Brown Norway F1 rats were obtained (Charles River Laboratories, Senneville, QC, Canada). All protocols were approved by the University of Guelph’s Animal Care Committee (AUP #4905) and followed guidelines from the Canadian Council on Animal Care. Rats were housed at 23°C in groups of two or three and given ad-libitum access to a Teklad global 18% protein rodent diet (Envigo, Huntington, Cambs., UK) and room-temperature water. As described in detail below, rats were subjected to 2 weeks of unilateral immobilization of the plantar flexors in a shortened position. Thrice weekly during the casting immobilization period, casts were removed and the plantar flexors were subjected to eccentric training, after which a new cast was immediately applied. Mechanical testing was performed 2-7 days prior to application of casts, then again 48 hours following the final training session, immediately following cast removal. Following the final mechanical testing session, rats were euthanized via isoflurane followed by CO2 asphyxiation and cervical dislocation. The hindlimbs were amputated and fixed in formalin for subsequent determination of soleus and medial gastrocnemius (MG) SSN. In accordance with previous studies ^20,37–40^, the left leg served as the experimental leg while the right leg served as an internal control.

### Data acquisition during mechanical testing and training

A 701C High-Powered, Bi-Phase Stimulator (Aurora Scientific, Aurora, ON, Canada) was used to evoke transcutaneous muscle stimulation via a custom-made electrode holder with steel galvanized pins situated transversely over the popliteal fossa and the calcaneal tendon ^5,6,33^. During piloting, we determined this stimulation setup produced similar values of maximum isometric tetanic torque across repeated testing sessions, and produced plantar flexion torque values consistent with values in previous literature using direct nerve stimulation ^41^. Torque, angle, and stimulus trigger data were sampled at 1000 Hz with a 605A Dynamic Muscle Data Acquisition and Analysis System (Aurora Scientific, Aurora, ON, Canada) with a live torque tracing during all training and mechanical data collection sessions.

### Mechanical testing

As described previously ^5^, rats were anesthetized with isoflurane and positioned supine on a heated platform (37°C) (Figure 1A). The left leg was shaved and fixed to a force transducer/length controller foot pedal via tape, with the knee immobilized at 90°. Each mechanical testing session began with determination of the optimal stimulation current for active plantar flexion torque (frequency = 100Hz, pulse duration = 0.1ms, train duration = 500ms) at an ankle angle of 90° (full plantar flexion = 180°) (Figure 1D), which was the current used throughout the remainder of the testing session. This stimulation current was confirmed to maximally activate the plantar flexors with minimal spread to the antagonist muscles as described previously ^5^. A 100Hz stimulation was then completed at an ankle angle of 70° (Figure 1E). Active torque was measured by subtracting the torque value at baseline (passive torque) immediately prior to stimulation from the total torque during stimulation ^42,43^. A passive torque-angle relationship was then constructed by recording passive torque following 5s of stress-relaxation at ankle angles of 100, 95, 90, 85, 80, 75, and 70° (Figure 1F). Isotonic (i.e., constant torque) contractions were then performed at load clamps equating to 30% and 40% of the maximum isometric torque at 70° (Figure 1G-H), in a randomized order, as we previously determined these loads to represent peak isotonic power for the rat plantar flexors ^5,33^. Peak angular velocity was recorded as the maximum time derivative of the angular displacement during the isotonic contraction ^5^, and power was recorded as the torque of the load clamp multiplied by peak angular velocity. Power and angular velocity were averaged between the 30% and 40% isotonic contractions for statistical analyses. Two minutes of rest separated each stimulation to minimize the development of muscle fatigue.

**Figure 1:**
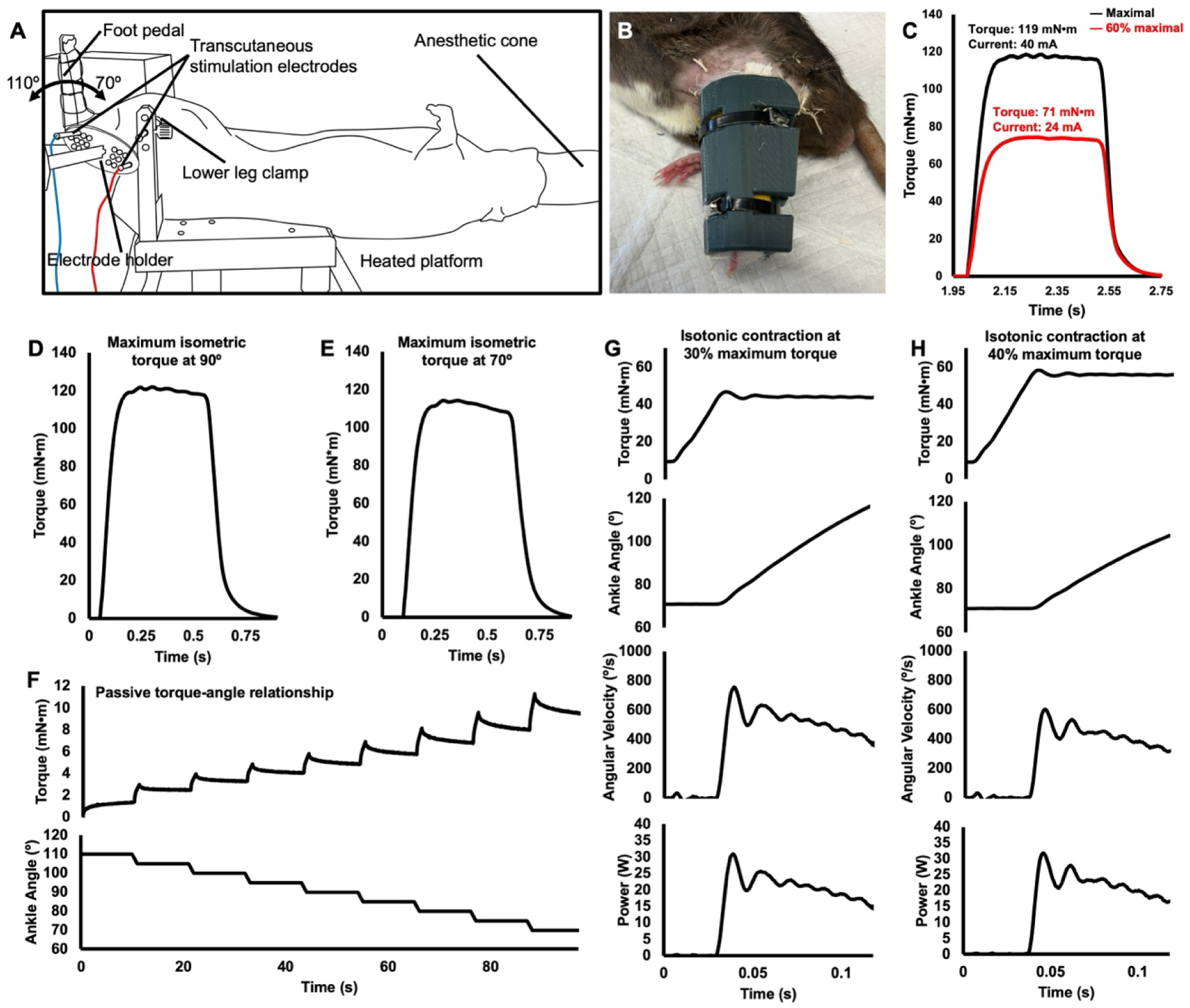
Experimental setup and representative traces for mechanical testing data from an old rat. **A.** Experimental setup for mechanical testing and eccentric training. The rat’s left foot was secured to a foot pedal and electrically stimulated to evoke plantar flexion contractions, with the knee fixed at 90°. The ankle had a 70° to 110° range of motion. **B.** Unilateral cast placing the ankle in full plantar flexion. **C.** Raw torque traces of a maximum isometric contraction and the corresponding 60% maximum isometric contraction. The stimulation current from the 60% maximum isometric contraction was used for the remainder of that training session during the eccentric contractions. **D.** Raw torque trace of a maximum isometric contraction at an ankle angle of 90°. **E.** Raw torque trace of a maximum isometric contraction at an ankle angle of 70°. **F.** Raw torque and ankle angle traces for construction of the passive torque-angle relationship. Passive torque was recorded immediately prior to each length change. **G-H**. Raw torque, ankle angle, angular velocity, and power traces for isotonic contractions from 70° to 110° with load clamps set to 30% and 40% of the maximum isometric torque. Peak power was recorded as torque multiplied by the peak angular velocity.

### Unilateral immobilization

Using gauze padding, vet wrap, and a 3D-printed brace and splint, the right hindlimb of each rat was casted in full plantarflexion for 2 weeks ^6^ (Figure 1B). The toes were left exposed to monitor for swelling ^40,44^. Casts were inspected daily and repaired/replaced as needed. All casts were replaced 1 week into the casting period following the 1 wk cast ultrasound measurements. Since rats were free to walk around while wearing their casts, this intervention merely promoted disuse of the immobilized muscles rather than complete unloading such as that achieved with hindlimb suspension ^45^.

### Isokinetic eccentric training

The submaximal eccentric training protocol was replicated directly from our previous study ^33^, in which we observed 8-11% increases in SSN and a 23% increase in peak isotonic power in old rats. Training lasted 2 weeks (i.e., during the casting intervention) and occurred 3 days/week (Monday, Wednesday, Friday).

At the start of each training session, the stimulus current was first adjusted to produce maximum isometric torque (pulse duration = 0.1ms, frequency = 100Hz, train duration = 500ms) at an ankle angle of 90°. The current was then adjusted to produce an isometric torque that as closely as possible matched 60% of maximum isometric torque (Figure 1C). Maximum isometric torque was recorded for each training session to track changes in muscle strength throughout the training period.

Each eccentric repetition consisted of 3 phases: 1) a 500-ms pre-activation at 110°; 2) active lengthening to 70°; and 3) 3 seconds of deactivation followed by a return to 110° at 20°/s. An additional 3 seconds of rest were provided before the next repetition. During week 1, rats completed 3 sets of 8 repetitions at 40°/s. In week 2, they completed 3 sets of 9 repetitions at 40°/s. Two minutes of rest were provided between each set.

### Serial sarcomere number determination

Following their final mechanical testing session, rats were sacrificed via isoflurane anesthetization followed by CO2 asphyxiation and cervical dislocation. The hindlimbs were amputated, skinned with all muscles overlying the plantar flexors removed, and fixed in 10% phosphate-buffered formalin with the ankle pinned at 90°. After fixation for 1-2 weeks, the soleus and MG were dissected off the lower leg, weighed, then re-submerged in formalin until the commencement of SSN estimations. To commence the process of SSN estimations, the muscles were rinsed with phosphate-buffered saline, then digested in 30% nitric acid for 6-8 h to remove connective tissue and allow for individual muscle fascicles to be teased out ^43,46^.

For each muscle, two fascicles were obtained from each of the proximal, middle, and distal regions (i.e., n = 6 fascicles total per muscle). SSN and FL values were averaged across these six fascicles for the reporting of data. Dissected fascicles were placed on glass microslides (VWR International, USA), then FLs were measured using ImageJ software (version 1.53f, National Institutes of Health, USA) from pictures captured by a level, tripod-mounted digital camera, with measurements calibrated to a ruler in plane with the fascicles. Sarcomere length (SL) measurements were taken at n = 6 different locations proximal to distal along each fascicle via laser diffraction (Coherent, Santa Clara, CA, USA) with a 5-mW diode laser (∼1 mm beam diameter, 635 nm wavelength) and custom LabVIEW program (Version 2011, National Instruments, Austin, TX, USA) ^47^, for a total of n = 36 SL measurements per muscle. For each fascicle, the six SL measurements were averaged to obtain a value of average SL. Given the laser diameter of ∼1 mm, one SL measurement itself represents an average of hundreds to thousands of SLs. Our total quantity of SL and FL measurements is consistent with previous studies ^42,43,46^. For each fascicle, the six SL measurements were averaged to obtain a value of average SL. Serial sarcomere number of each fascicle was calculated as:

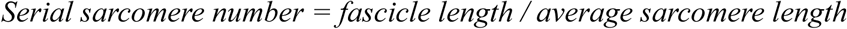

### Determination of physiological cross-sectional area

To gain further insight on changes in muscle contractile tissue in parallel, we calculated physiological cross-sectional area (PCSA; in cm^2^) using the equation ^42,48^:

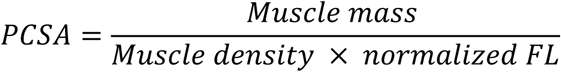

Muscle density was assumed to be 1.112 g/cm^3 48^. Normalized FL was calculated using FL of dissected fascicles and measured SL in the equation ^48^:

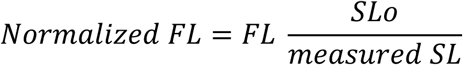

SLo represents optimal SL of rat muscle at rest, assumed to be ∼2.7 μm based on previous literature ^42,49^. Maximum isometric torque and peak isotonic power normalized to the combined PCSA of the soleus and MG were also recorded, with post-training values normalized to trained muscle PCSA, and pre-training values normalized to untrained muscle PCSA.

### Statistical analyses

All statistical analyses were performed in SPSS Statistics Premium version 28. Normality of data was confirmed using Shapiro-Wilk tests. Differences in SSN, SL, FL, muscle wet weight, and PCSA between the trained and untrained leg were assessed by two-way repeated measures analysis of variance (ANOVA) (Age [young, old] × Cast [un-casted leg, casted leg]). Differences in peak power were also assessed via two-way repeated measures ANOVA (Age [young, old] × Cast [pre-cast, post-cast]). Three-way repeated measures ANOVAs assessed training-induced changes in maximum isometric torque (Age [young, old] × Cast [pre-cast, post-cast] × Angle [90°, 70°]) and passive torque (Age [young, old] × Cast [pre-cast, post-cast] × Angle [110°-70°]). Three-way ANOVA was used in these cases to allow us to investigate differences in the overall shapes of the active and passive torque-angle relationships. Lastly, to investigate time course changes in maximum isometric torque at 90° across each day of training, we used a two-way repeated measures ANOVA (Age [young, old] × Training day [pre-cast, training day 1, 2, 3, 4, 5, 6, post-cast]). A Greenhouse-Geisser correction for sphericity was applied for all ANOVAs and a Sidak correction was applied to all pairwise comparisons. Significance was set at α=0.05.

## Results

### Changes in muscle mass and physiological cross-sectional area

For soleus wet weight, there was an effect of cast (F(1,18)=154.466, *P*<0.001) but no effect of age (F(1,18)=1.553, *P*=0.229) such that wet weight was 40% lower in the casted compared to control soleus, with no age-related difference (Figure 2A). Similarly, for soleus PCSA, there was an effect of cast (F(1,18)=25.204, *P*<0.001) but no effect of age (F(1,18)=0.021, *P*=0.886) nor an interaction (F1,18)=0.470, *P*=0.502). Thus, PCSA of the casted soleus was 20% lower than that of the control soleus, with no age-related difference (Figure 2B).

**Figure 2:**
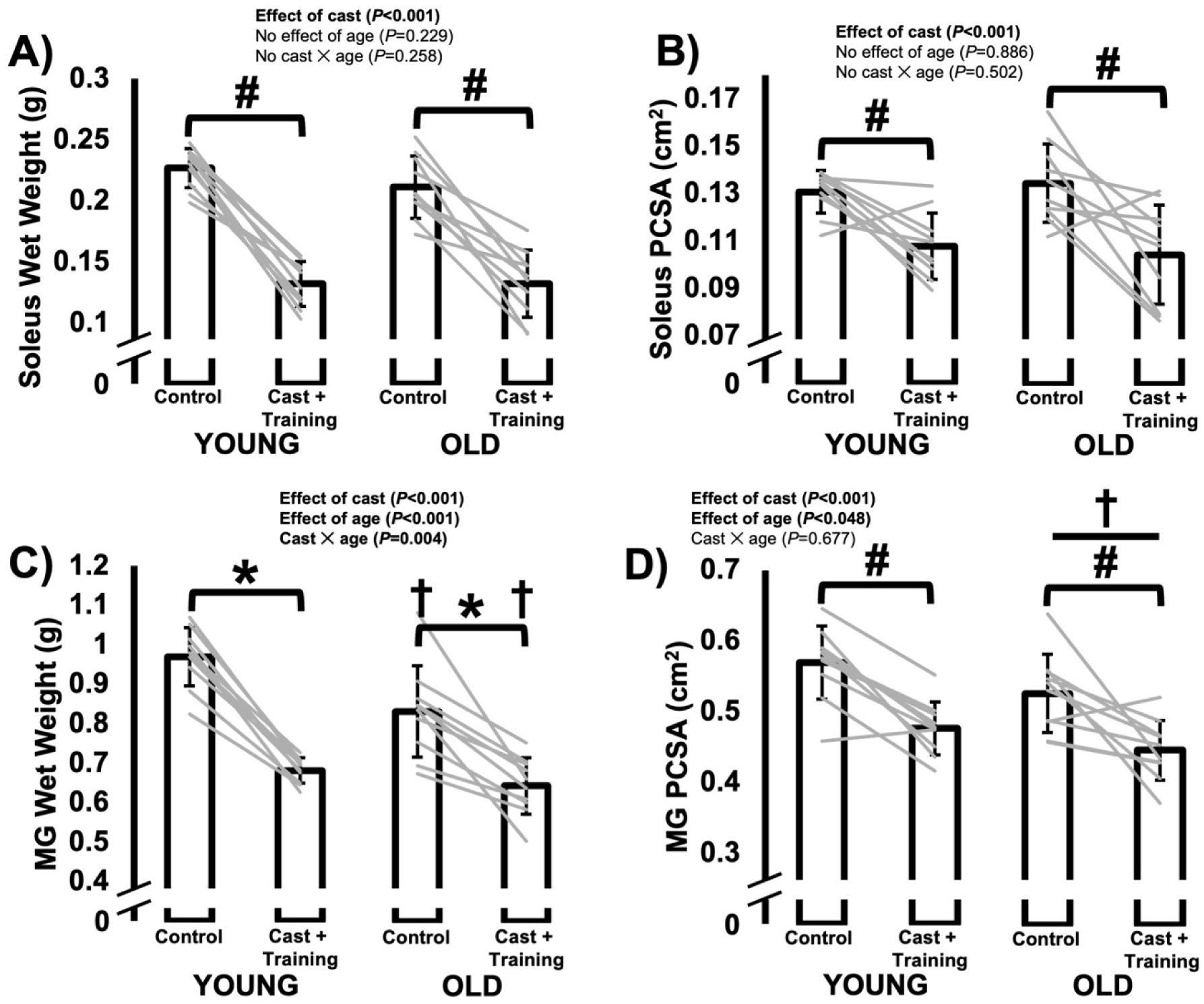
Differences in muscle wet weight and physiological cross-sectional area (PCSA) of the soleus (**A-B**) and medial gastrocnemius (MG; **C-D**) between the control leg vs. the leg that was casted with intermittent submaximal eccentric training, and between old vs. young rats. A, B, and D: #Effect of cast with data from young and old rats combined. †Effect of age with data from control and casted combined (*P*<0.05). C: *Difference between control and casted (*P*<0.05). **†**Difference between young and old rats (*P*<0.05).

For MG wet weight, there was a cast × age interaction (F(1,18)=10.601, *P*=0.004). Old rats had lower MG wet weight than young in both the control (−20%, *P*<0.001) and casted leg (−11%, *P*=0.004). Both age groups exhibited lower MG wet weight in the casted leg than the control leg (young: −30%, *P*<0.001; old: −22%, *P*<0.001) (Figure 2C). For MG PCSA (Figure 2D) there were effects of cast (F(1,18)=36.832, *P*<0.001) and age (F(1,18)=4.511, *P*=0.048) but no interaction (F(1,18)=0.179, *P*=0.677). MG PCSA was 6% smaller in old than young rats and 16% smaller in the casted than the control leg.

### Changes in muscle fascicle length driven by serial sarcomere number adaptations

For FL as measured at a 90° ankle angle in both the soleus and MG (Figure 3A-B), there were effects of age (soleus: F(1,18)=6.351, *P*=0.021; MG: F(1,18)=19.776, *P*<0.001) and cast (soleus: F(1,18)=126.945, *P*<0.001; MG: F(1,18)=54.480, *P*<0.001), but no interaction (soleus: F(1,18)=0.576, *P*=0.458; MG: F(1,18)=3.015, *P*=0.100) such that FL was shorter in old than young rats (soleus: –5%; MG: –8%) and shorter in the casted than the control leg (soleus: –22%; MG: – 12%).

**Figure 3:**
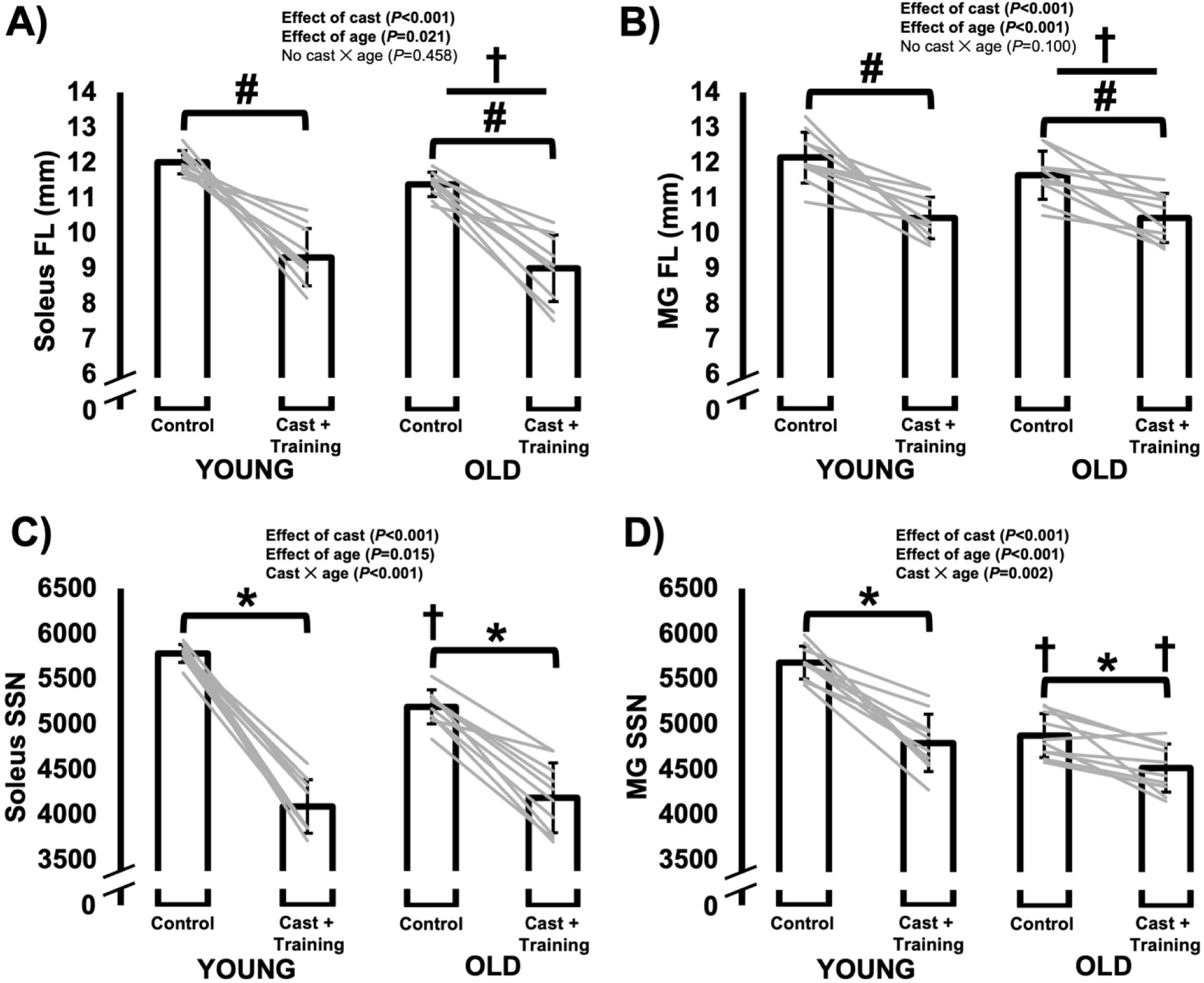
Differences in muscle fascicle length (FL) and serial sarcomere number (SSN) of the soleus (**A, C**) and medial gastrocnemius (MG; **B, D**) between the control leg vs. the leg that was casted with intermittent submaximal eccentric training, and between old vs. young rats. A-B: #Effect of cast with data from young and old rats combined. †Effect of age with data from control and casted combined (*P*<0.05). C-D: *Difference between control and casted (*P*<0.05). **†**Difference between young and old rats (*P*<0.05).

The SSN data showed that the above age-related and cast-related reductions in FL and overstretched SLs (see data in Supplemental Figure S1) were driven by reductions in SSN (Figure 3E-F). For both the soleus and MG, there were cast × age interactions (soleus: F(1,18)=24.000, *P*<0.001; MG: F(1,18)=13.524; *P*=0.002). In the soleus, SSN was lower in the casted than the control leg for old (–19%, *P*<0.001) and young rats (–29%, *P*<0.001). The greater magnitude decrease for young rats corresponded to a lower SSN in old than young rats for the control leg (– 10%, *P*<0.001) but not the casted leg (*P*=0.544). The MG also exhibited cast-related losses of SSN in both old (–7%, *P*=0.003) and young rats (–16%, *P*<0.001), although at a smaller magnitude than in the soleus. Old rats had a lower MG SSN than young rats in both the control (–14%, *P*<0.001) and casted leg (–6%, *P*=0.048).

### Changes in maximum isometric torque as a function of joint angle

For maximum isometric torque, there was a cast × joint angle × age interaction (F(1,18)=421.745, *P*<0.001). Old rats produced 36-44% less isometric torque than young rats at ankle angles corresponding to both neutral (90°) and stretched (70°) muscle lengths pre- and post-cast (all *P*<0.001) (Figure 4A). Furthermore, both old and young rats experienced decreases in torque at both joint angles from pre- to post-cast, with the decreases at 70° noticeably larger in magnitude (old 90°: −37%; old 70°: −44%; young 90°: −37%; young 70°: −46% all *P*<0.001) (Figure 4A). Therefore, intermittent submaximal eccentric training during casting did not prevent the loss of maximum isometric strength, with torque at a more stretched muscle length especially affected. This greater effect on torque at 70° was most noticeable in young rats, because pre-training, torque at 70° was greater than torque at 90° (*P*<0.001), whereas post-training, torque at 90° was instead greater than torque at 70° (*P*<0.001), indicating a leftward shift in the torque-angle relationship (Figure 4A) towards shorter muscle lengths. For old rats, torque was greater at 90° than 70° both pre- and post-training (both *P*<0.001).

**Figure 4:**
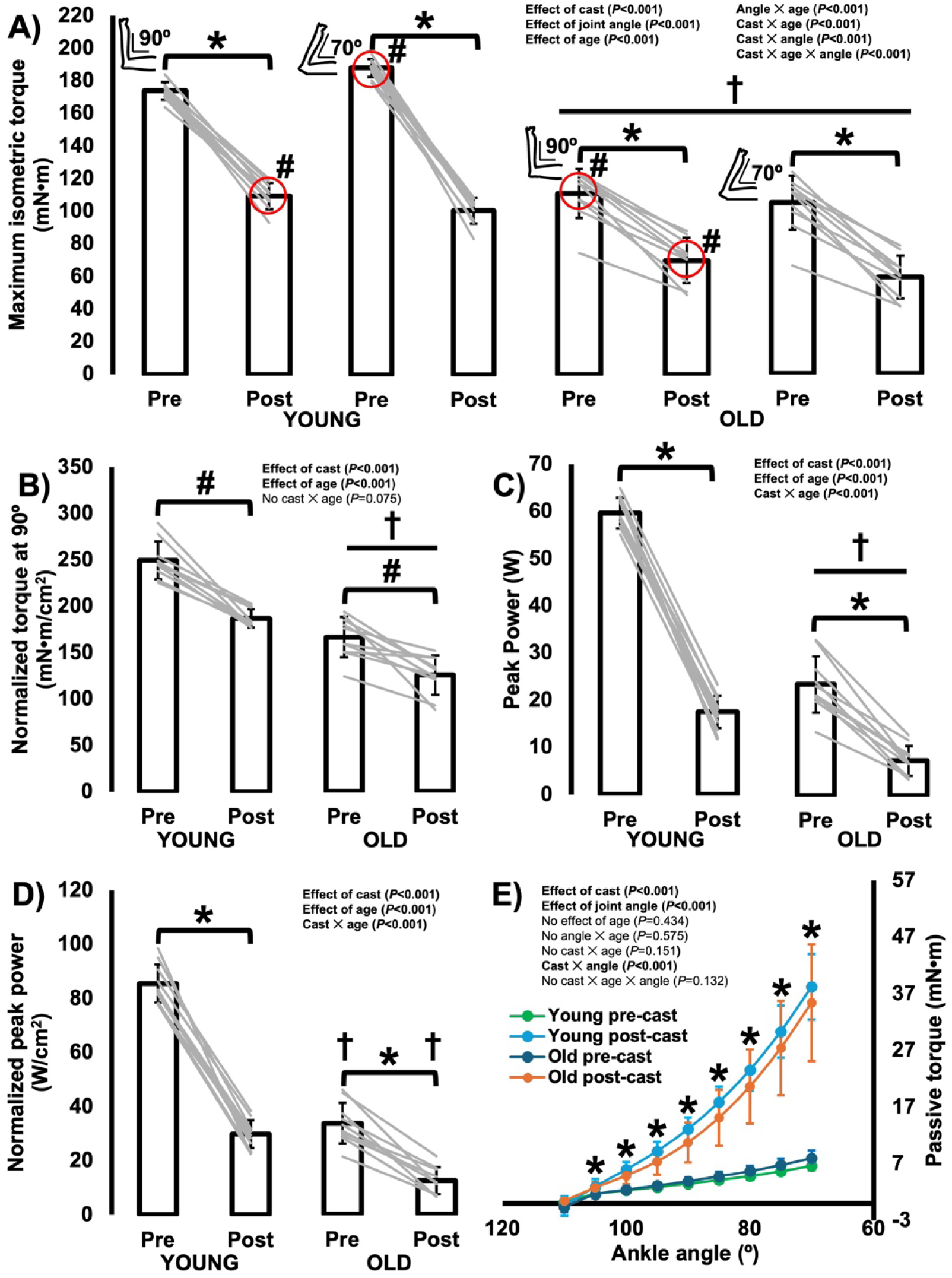
**A.** Differences in maximum isometric torque as a function of joint angle from pre- to post-cast and between young and old rats. *Difference from pre- to post-cast (*P*<0.05). **†**Difference between young and old rats (*P*<0.05). #Difference between ankle angles of 90° and 70° (*P*<0.05), with the red circle indicating which angle produced more optimal torque at pre- and post-training. **B.** Differences in maximum isometric torque at 90° normalized to the combined physiological cross-sectional area of the soleus and medial gastrocnemius from pre- to post-cast and between young and old rats. #Effect of cast with data from young and old rats combined (*P*<0.05). **†**Effect of age with data from control and casted combined (*P*<0.05). **C.** Differences in peak isotonic power from pre- to post-cast and between young and old rats. *Difference from pre- to post-cast (*P*<0.05). **†**Difference between young and old rats (*P*<0.05). **D.** Differences in peak power normalized to the combined physiological cross-sectional area of the soleus and medial gastrocnemius from pre- to post-cast and between young and old rats. *Difference from pre- to post-cast (*P*<0.05). **†**Difference between young and old rats (*P*<0.05). **E.** Differences in passive torque as a function of joint angle from pre- to post-cast and between young and old rats. *Difference between pre- and post-cast (*P*<0.05) with data from young and old rats pooled, as there was no interaction with age.

After normalizing maximum isometric torque at 90° to the combined PCSA of the soleus and MG, there was still an effect of age (F(1,18)=133.933, *P*<0.001), with old rats producing 33% lower normalized torque than young (Figure 4B). There was also an effect of cast (F(1,18)=80.873, *P*<0.001) but no cast × age interaction (F(1,18)=3.576, *P*=0.075) for normalized torque, with normalized torque decreasing 25% from pre- to post-cast. These losses of normalized torque indicate losses of muscle quality both with aging and from pre- to post-cast.

### Time-course losses of maximum isometric torque at 90° throughout the casting/intermittent training period

For maximum isometric torque at 90° across the 6 days of intermittent training during casting, there was a training day × age interaction (F(2.24,40.279)=12.886, *P*<0.001), indicating the time course of torque loss during casting differed between old and young rats. Old rats exhibited torque loss starting on the second day of training (*P*<0.001 compared to pre-cast and day 1 of training), then continued to lose torque up to the fourth day of training (*P<*0.001-0.018 compared to all prior days), after which the loss of torque plateaued because torque in the final three days of training did not differ compared to each other and compared to post-cast (*P*=0.062-1.00) (Figure 5). Young rats, conversely, underwent a progressive loss of maximum isometric torque throughout the whole casting period starting after the first day of training, with torque at each subsequent day of training and post-cast differing from all other days (*P*<0.001-0.002) (Figure 5).

**Figure 5:**
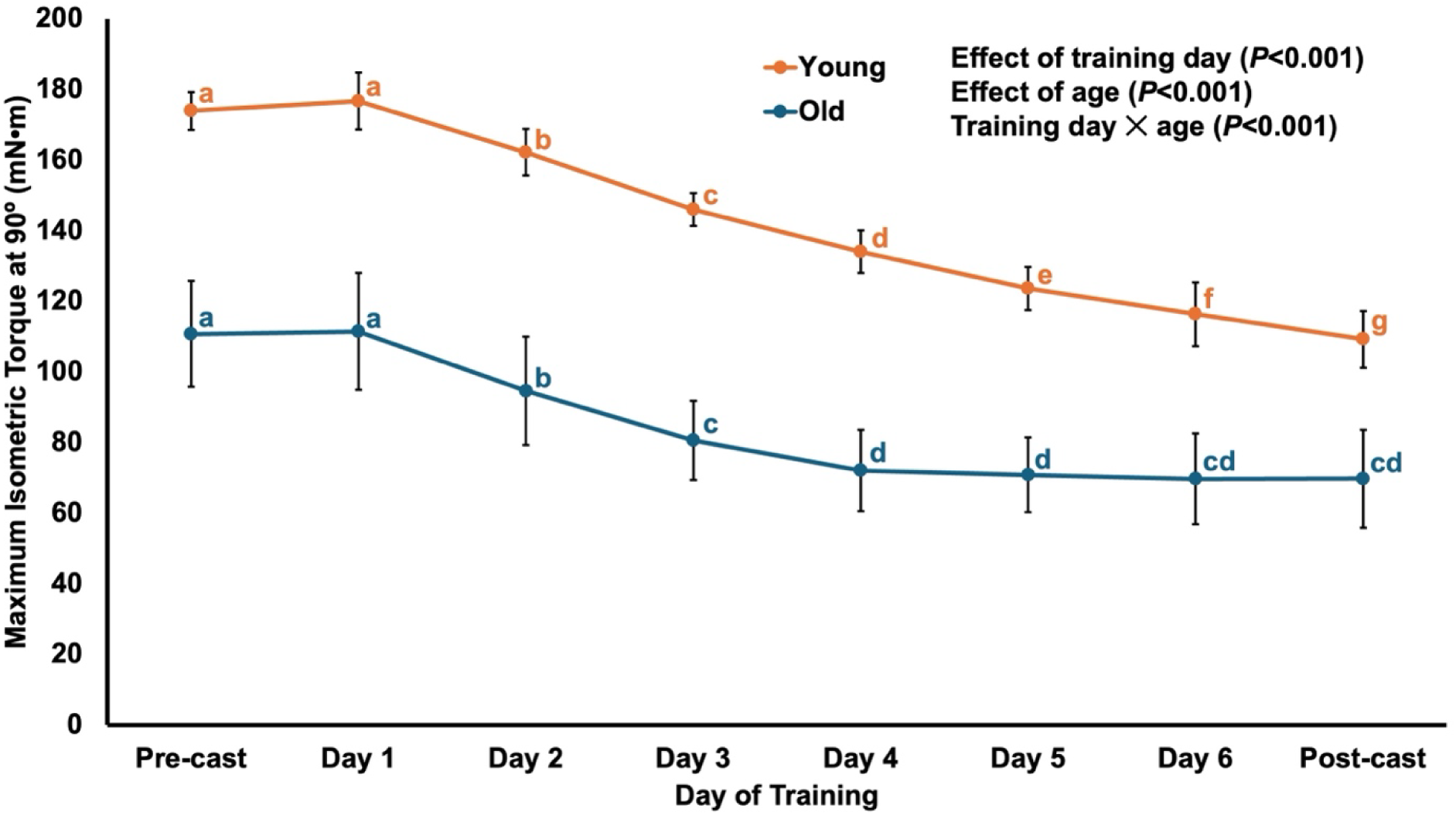
Changes in maximum isometric torque at 90° throughout the 2-week casting with intermittent training period. Training began two days after application of casts and occurred three days per week (Monday, Wednesday, Friday), amounting to 6 training sessions in total. Casts were removed and post-cast testing was completed two days following the final training session. Different letters denote a significant difference between time points (*P*<0.05).

### Changes in peak isotonic power

For peak isotonic power, there was a cast × age interaction (F(1,18)=156.35, *P*<0.001). Old rats were 58-60% less powerful than young rats pre- and post-training (both *P*<0.001) (Figure 4C). Old and young rats also both experienced losses of power from pre- to post-cast, with a 69% loss in old rats and a 71% loss in young rats (both *P*<0.001) (Figure 4C). These losses of power seemed to be due to both the losses of torque and angular velocity, because peak angular velocity during the isotonic contractions also had a cast × age interaction (F(1,18)=11.885, *P*=0.003) decreasing 47% and 46% from pre- to post-cast in old and young rats, respectively (both *P*<0.001). Collectively, intermittent submaximal eccentric training during casting did not prevent the losses of torque, velocity, or power generation, with power undergoing the largest decrease.

After normalizing peak isotonic power to the combined PCSA of the soleus and MG (Figure 4D), there was still a cast × age interaction (F(1,18)=90.222, *P*<0.001), with lower normalized power in old compared to young rats (pre-training: –59%, *P*<0.001; post-training: – 56%, *P*<0.001), and post-training compared to pre-training (young: –65%, *P*<0.001; old: –63%, *P*<0.001). Therefore, the loss of PCSA only accounted for a small portion of the loss of peak power from pre- to post-cast.

### Changes in passive torque as a function of joint angle

For passive torque, there was a cast × joint angle interaction (F(1.03,18.57)=251.54, *P*<0.001), and no effect of age (F(1,18)=0.639, *P*=0.434), indicating passive torque changed differently across joint angles from pre- to post-cast, but was consistent between old and young rats. As expected, both pre- and post-cast, passive torque increased with increasing muscle stretch (all comparisons (*P*<0.001). From pre- to post-cast, passive torque underwent increases at all joint angles (all *P*<0.001) other than 110° (*P*=0.645), with the greatest increases of 404% at 70° (i.e., the most stretched joint angle tested) (Figure 4E). Therefore, intermittent submaximal eccentric training during casting did not prevent the elevation of passive torque that often occurs following casting in a shortened position.

## Discussion

Based on our previous observations of serial sarcomerogenesis and improved muscle mechanical performance with submaximal eccentric training in old rats ^33^, this study aimed to prevent the loss of SSN during a period of immobilization-induced disuse (i.e. casting in a shortened muscle position) via intermittent submaximal eccentric training. We observed losses of muscle wet weight, SSN, PCSA, and maximal strength and power production in both old and young rats. Thus, submaximal eccentric training 3 days/week did not counteract the effects of casting, and this was not age dependent.

### Influence of aging- and casting-induced changes in muscle morphology on mechanical function

Consistent with previous reports, old rats exhibited lower muscle wet weights ^41,50–52^ and shorter FLs driven by a reduced SSN ^3–6^ as compared to young rats. This loss of SSN likely contributed to old rats producing optimal torque at a more shortened muscle length (90° ankle angle) than young rats (70° ankle angle) (Figure 4A), and producing lower peak isotonic power (Figure 4C) due to reduced torque production throughout the range of motion ^18^. These observations align with our previous work ^5,6,33^ and studies from other groups as well ^41,51,53^. Notably, there was only an age-related loss of PCSA in the MG and not the soleus. Our previous study also observed no age-related loss of soleus PCSA ^6^, and given the weaker torque-production capacity in the old compared to young rats, indicates reduced muscle quality ^54,55^. Reduced muscle quality underscores age-related changes to the intrinsic contractile machinery such as reduced myosin protein content or oxidation of myosin, which reduces crossbridge formation and force production ^56,57^.

Following 2 weeks of casting with intermittent submaximal eccentric training, we observed losses of muscle wet weight in the soleus and MG, which seemed to be due to losses of contractile tissue both in series (SSN) and in parallel (PCSA) (Figures 2 and 3). These muscle morphological changes likely drove the losses of maximum isometric torque and peak isotonic power (Figure 4). The losses of SSN, PCSA, and muscle wet weight were greater for the soleus than the MG, likely because our casts only fully immobilized the ankle while not limiting mobility at the knee. The MG crosses both the knee and ankle whereas the soleus only crosses the ankle, therefore, there was likely some loading of the MG at the knee that allowed better maintenance of MG morphology compared to the soleus.

In young rats, there was also a shift in the optimal angle for torque production from a more stretched muscle length pre-training to a more neutral muscle length post-training (Figure 4A), which is consistent with the loss of SSN due to shifting of the optimal muscle length for actin-myosin overlap ^6,20,38,58^. As power is the dot product of force and velocity, the loss of torque generating capacity likely contributed to the loss of peak isotonic power. However, since normalizing power to PCSA only slightly reduced the cast-related power loss (from ∼70% to ∼65%), other factors besides the loss of force-production capacity could have contributed, notably two main factors. First, the loss of muscle quality (i.e., maximum torque normalized to PCSA; Figure 4B) could have contributed by reducing intrinsic crossbridge function. Second, the loss of SSN could have contributed to power loss by narrowing the torque-angle relationship’s plateau region and reducing torque-production capacity throughout the joint range of motion ^18^. These findings highlight the functional implications of SSN loss and the importance of SSN maintenance throughout the lifespan for muscle mechanical performance.

### No benefit of casting with intermittent submaximal eccentric training compared to casting alone

Many of the changes from pre- to post-cast we observed here are similar in magnitude to those observed in our previous study following 2 weeks of casting without intermittent training ^6^, as displayed in Supplemental Figure S2. In fact, it appears that if anything, intermittent submaximal eccentric training during casting was more detrimental than casting alone, as maximum passive torque increased ∼3-fold more (+343-478% vs. +68-180%) and muscle wet weight decreased more (37-42% vs. –23-25%) with intermittent eccentric training during casting (Supplemental Figure S2). Thus, not only did intermittent submaximal eccentric training not prevent the effects of casting in a shortened position, the two together may have had a compounded negative effect on muscle function. While histological assessment of the extracellular matrix was beyond the scope of the present study, the greater increase in passive torque compared to our previous study may be indicative of greater fibrous connective tissue deposition ^59–61^. It is possible that overstretched sarcomeres due to the loss of SSN following casting could have contributed to the elevated passive torque, however, we only observed a casting-induced increase in SL as measured at 90° in young rats, with no change in old rats (Supplemental Figure S1). Furthermore, a similar increase in SL was also observed in our previous study of casting alone ^6^ alongside the smaller casting-induced increase in passive torque. Thus, it seems more likely that changes in extracellular matrix morphology contributed to the elevated passive torque here.

While speculative, it is possible some muscle damage was incurred during the eccentric contractions that led to a worsened effect compared to casting alone. Shorter muscles (i.e., with fewer sarcomeres in series) can undergo more damage during eccentric contractions for a given magnitude of joint rotation because the stretch imposed on individual sarcomeres would be greater ^62,63^. Hence, it is possible that as the muscle lost SSN from casting in a shortened position, it became increasingly more susceptible to eccentric contraction-induced damage. In support of this theory, previous studies have shown that the effects of eccentric-induced muscle damage are amplified if preceded by immobilization ^64^, and that immobilization immediately following eccentric exercise may also prevent recovery from muscle damage ^65^. Further work is needed to elucidate the role of muscle damage in intermittent submaximal eccentric training during casting.

### Comparison to other intermittent training interventions employed during immobilization

We can also look to previous studies that employed intermittent passive stretching during casting to gain insight into why our intermittent eccentric training did not prevent the deleterious effects of casting on muscle. Williams (1990) found that exposing the mouse soleus to 30 minutes to 2 hours/day of passive stretching prevented the loss of SSN during casting in a shortened position. They even observed a 10% increase in soleus SSN compared to controls in the legs stretched for 2 hours/day during casting. Passive stretch training alone (i.e., in non-immobilized muscle), however, does not always result in increased SSN depending on the duration and intensity ^29,30,66,67^. It seems in general that shorter duration or less frequent passive stretch training has minimal success for increasing SSN. It follows that Gomes et al. (2004) and Coutinho et al. (2004) did not prevent SSN loss during casting with intermittent passive stretch training 1 or 3 days/week, respectively in rats. There was reason to believe that intermittent submaximal eccentric training 3 days/week would have been more successful than passive stretch training at preventing SSN loss during casting. While eccentric training interventions have also shown varying levels of success for increasing SSN ^5,42,43,46,68,69^, the most robust increase in SSN following eccentric training was in our previous study, with 8-11% increases in soleus and MG SSN following 4 weeks of submaximal isokinetic eccentric training 3 days/week in old rats ^33^. Furthermore, Van Dyke et al. (2012) found that stretch and contraction combined prevented the loss of SSN following tenotomy while passive stretching did not. Collectively, one might expect the conditions of muscle stretch and activation provided by eccentric training to have better success than passive stretch training for preventing SSN loss during casting. However, similar to the findings of Coutinho et al. (2004) with passive stretch training 3 days/week, it is possible that eccentric training for ∼10 min/day 3 days/week in the present study was not enough to be the dominant stimulus imposed on the muscle compared to casting in a shortened position.

Given the use of transcutaneous electrical stimulation for the present study’s submaximal eccentric training, comparisons can also be drawn to studies that employed “neuromuscular electrical stimulation” (NMES) to counteract the effects of immobilization on skeletal muscle. In NMES trials in humans, transcutaneous electrical stimulation is typically applied to the muscle as isometric contractions in 30- to 60-minute sessions once or twice daily, using a stimulation intensity set by the tolerance of the patient ^71–76^. NMES is well-recognized for preventing the reduction in myofibrillar protein synthesis and whole-muscle and myofiber cross-sectional area during immobilization, but is less successful at limiting the loss of muscle strength ^77^. Similar isometric electrical stimulation interventions in rats have also prevented myofiber atrophy and fibrosis during immobilization ^78–80^. These studies on rats used 15-20 minutes of tetanic electrical stimulation as few as 3 days/week at a stimulation intensity of up to 60% of the maximum torque development. These training parameters are similar to what was employed in the present study, other than that the present study used eccentric rather than isometric contractions, suggesting tentatively that isometric contractions may be more effective than eccentric contractions during immobilization.

To argue against the above perspective on isometric contractions, however, studies of conventional resistance training (i.e., concentric and eccentric contractions together) during bed rest in humans have also shown success for limiting losses of muscle function. Both plantar flexor ^81^ and leg press ^82^ resistance training every other day during 2 weeks of bed rest at up to 85% of the 1-repetition maximum prevented losses of 1-repetition maximum weight, concentric and eccentric strength, power production, and myofiber cross-sectional area. Even more impressive, squat and calf press resistance training every 3 days during 90 days of bed rest preserved muscle volume and force and power production ^83^. Considering the present study’s profound negative effects following casting with intermittent eccentric training, future studies are needed to elucidate optimal training stimuli during forced muscle disuse, especially for older individuals.

## Conclusion

Older adult humans are prone to falls and injuries that could result in extended periods of hospitalization, bed rest and muscle disuse, which can result in the loss of SSN on top of that already lost with aging. Thus, it is prudent to discover interventions to prevent the losses of SSN and associated aspects of mechanical function during immobilization in aged muscle. We sought to determine whether submaximal eccentric training could be one of such interventions. While we observed previously that submaximal eccentric training induced serial sarcomerogenesis and improved dynamic performance in muscle of old rats, we found here that this training provided no positive benefits to muscle morphology or function when employed during immobilization. Instead, large losses of muscle mass occurred due to losses of both SSN and PCSA, leading to large reductions in maximal torque and power generating capacity in both old and young rats.

## Acknowledgements

This project was supported by the Natural Sciences and Engineering Research Council of Canada (NSERC). The animals were obtained from the National Institute on Aging (NIA) aged rodent colonies.

## Conflict of interest statement

No conflicts of interest, financial or otherwise, are declared by the authors.

## Ethics statement

Approval was given by the University of Guelph’s Animal Care Committee and all protocols followed CCAC guidelines (AUP #4905).

## Data availability

All data generated or analysed during the study are available from the corresponding author upon request.

## Funding

This project was supported by the Natural Sciences and Engineering Research Council of Canada (NSERC), grant number RGPIN-2024-03782.

## Author contributions

A.H. and G.A.P. conceived and designed research; A.H. and E.V. performed experiments; A.H. analyzed data; A.H., E.V., and G.A.P. interpreted results of experiments; A.H. prepared figures; A.H. and G.A.P. drafted manuscript; A.H., E.V., and G.A.P. edited and revised manuscript; A.H., E.V., and G.A.P. approved final version of manuscript.

## Supplemental Figure S1

For soleus SL as measured at 90°, there was a cast × age interaction (F(1,18)=18.523, *P*<0.001). In young rats, SL was 10% longer in the casted than the control leg (*P*<0.001), whereas in old rats SL did not differ between legs (*P*=0.319). SL was 5% longer in old than young rats (*P*=0.002) for the control soleus but 6% shorter in old than young rats (*P*=0.012) for the casted soleus.

While there was also a cast × age interaction (F(1,18)=8.132, *P*=0.011) for MG SL as measured at 90°, it showed different patterns than for the soleus. In young rats, MG SL did not differ between legs (*P*=0.087), whereas in old rats MG SL of the casted leg was 3% shorter than the control leg (*P*=0.039). MG SL was 5% longer in old than young rats for the control leg (*P*=0.009) but not different between ages for the casted leg (*P*=0.990).

**Supplemental Figure S1:**
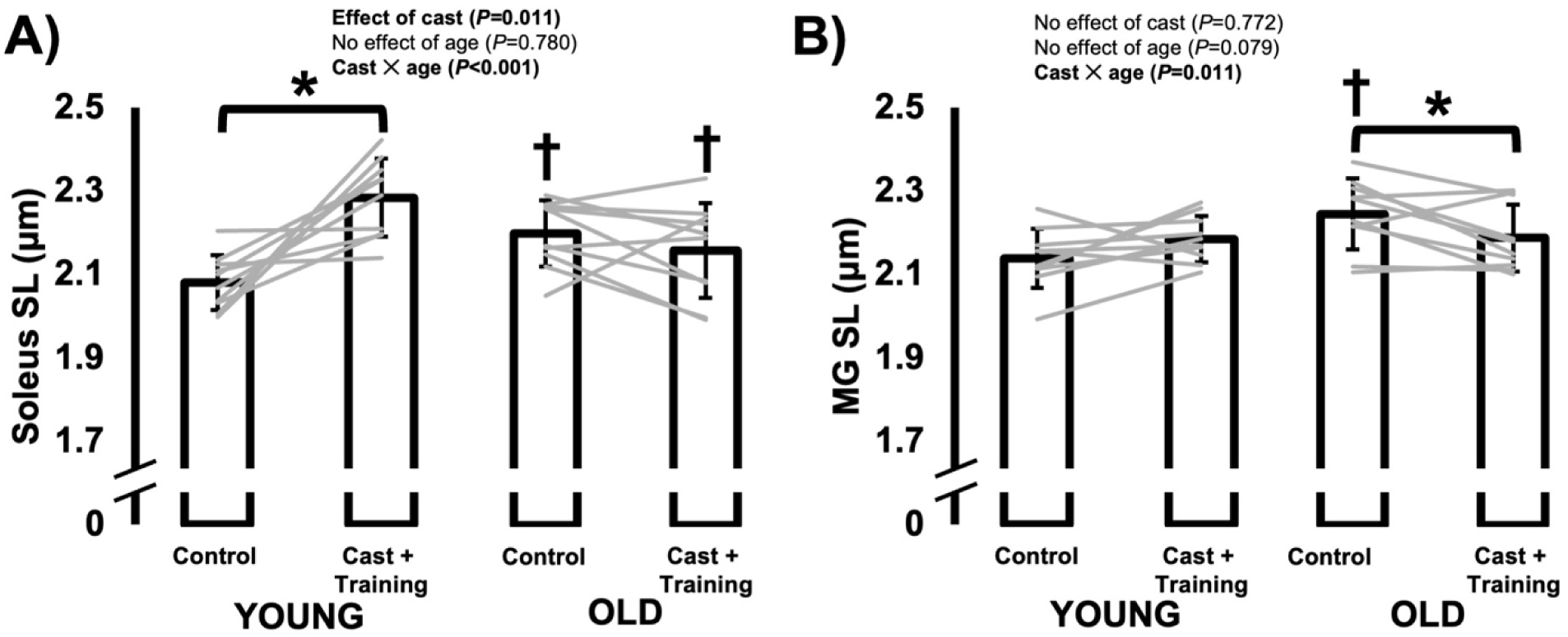
Differences in sarcomere length (SL) of the soleus (**A**) and medial gastrocnemius (MG; **B**) between the control leg vs. the leg that was casted with intermittent submaximal eccentric training, and between old vs. young rats. *Difference between control and casted (P<0.05). **†**Difference between young and old rats (P<0.05).

## Supplemental Figure S2

**Supplemental Figure S2:**
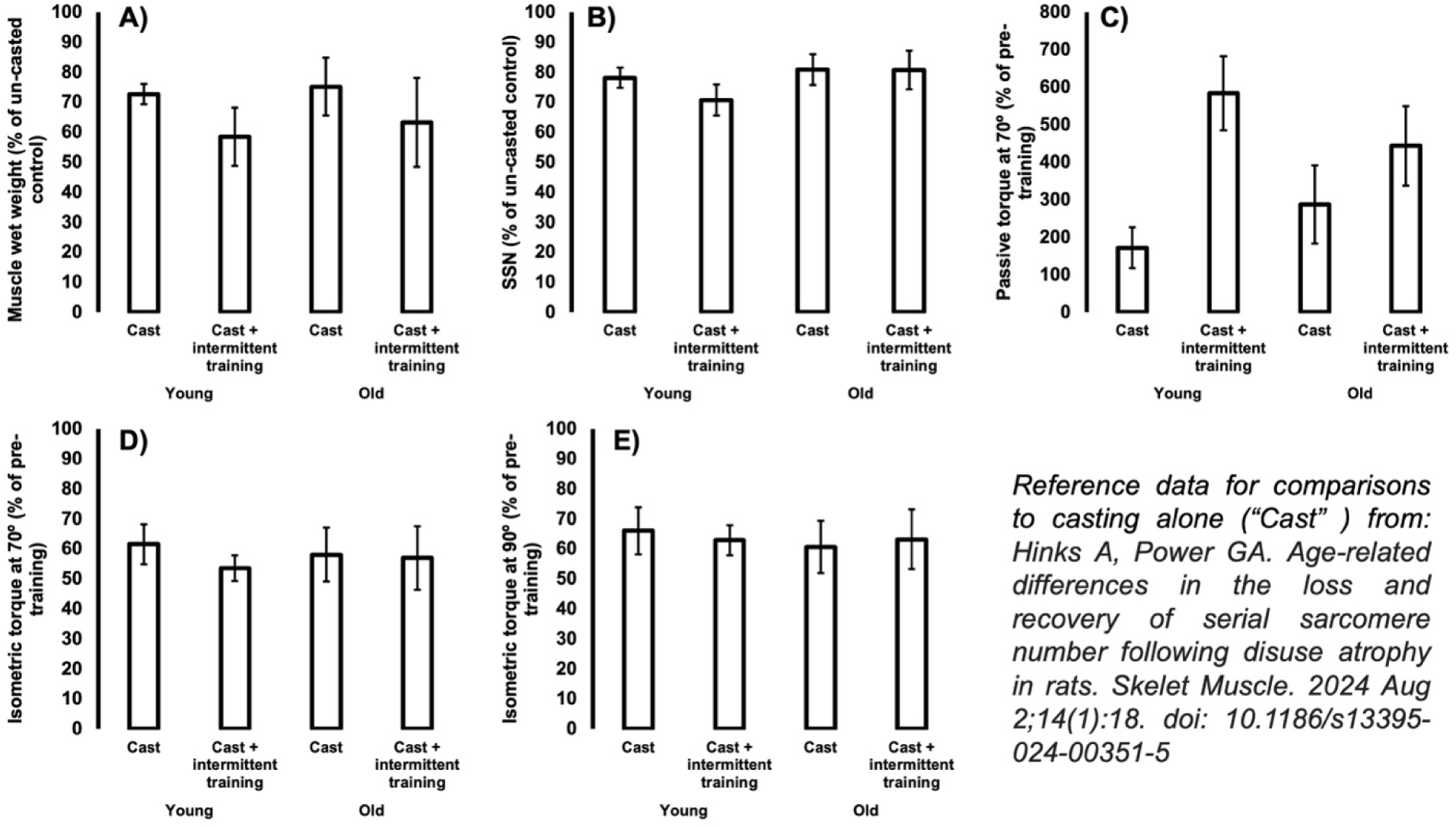
Comparisons of cast-induced changes in muscle wet weight (**A**), serial sarcomere number (SSN) (**B**), maximum passive torque (**C**), and maximum isometric torque at ankle angles of 70° (**D**) and 90° (**E**) between the present study’s casting with intermittent submaximal eccentric training (“Cast + intermittent training”) and our previous study of just casting (“Cast”) ^6^.

## Notes

### Competing Interest Statement

The authors have declared no competing interest.

